# An unsupervised feature extraction and selection strategy for identifying epithelial-mesenchymal transition state metrics in breast cancer and melanoma

**DOI:** 10.1101/865139

**Authors:** David J. Klinke, Arezo Torang

## Abstract

Digital cytometry is opening up new avenues to better understand the heterogeneous cell types present within the tumor microenvironment. While the focus is towards elucidating immune and stromal cells as clinical correlates, there is still a need to better understand how a change in tumor cell phenotype, such as the epithelial-mesenchymal transition, influences the immune contexture. To complement existing digital cytometry methods, our objective was to develop an unsupervised gene signature capturing a change in differentiation state that is tailored to the specific cellular context of breast cancer and melanoma, as a illustrative example. Towards this aim, we used principal component analysis coupled with resampling to develop unsupervised gene expression-based state metrics specific for the cellular context that characterize the state of cellular differentiation within an epithelial to mesenchymal-like state space and independently correlate with metastatic potential. First developed using cell line data, the orthogonal state metrics were refined to exclude the contributions of normal fibroblasts and to provide tissue-level state estimates based on bulk tissue RNA-seq measures. The resulting gene expression-based metrics for differentiation state aim to inform a more holistic view of how the malignant cell phenotype influences the immune contexture within the tumor microenvironment.

## Introduction

Tissues are comprised of a diverse set of different cell types that help maintain homeostasis. Oncogenesis is associated with a shift in the cellular composition of a tissue that can be revealed with increasing confidence through direct measurement, such as scRNA-seq, or using digital methods to deconvolute bulk tissue samples (1). Given the correlation with response to immunotherapies, the current focus has been on quantifying immune cell types present within the tumor microenvironment (2, 3). There is also an increasing appreciation for characterizing the heterogeneity among malignant cells that may arise in the same anatomical location (4, 5). Given our interest in understanding functional heterogeneity of malignant cells that originate within a particular anatomical organ rather than uncertainty in etiology, we will focus on breast cancer and melanoma as Li et al. show that melanoma and breast cancer cell lines seem to cluster most uniformly while other cell lines defined by anatomical origin seem to have a more heterogeneous composition (6).

While the tumor cells that arise in the skin and breast seem to be most similar, patient treatment strategies and outcomes can be diverse. Initial treatment strategies are guided by specific molecular alterations that can be targeted by drugs: aromatase inhibitors for ER-positive breast cancer, anti-HER2 antibodies for HER2-positive breast cancer, or small molecule inhibitors for BRAF V600E-positive or C-KIT-positive melanoma (7, 8). However, dissemination of primary tumors to vital organs like liver, brain, and lungs is a key limiter for patient survival in breast cancer and melanoma. Specifically, the 5-year survival rate for patients with localized disease versus distant metastases drops from 98% to 23% and from 99% to 27% for melanoma and breast cancer, respectively (9). In contrast, patient survival for tumors that originate in vital organs is limited by the degree to which malignant cells locally disrupt organ function. Thus the importance of distal dissemination in determining patient outcomes can vary based on the tissue of origin.

The distal dissemination and growth of malignant cells - metastasis - is a complex process thought to involve dynamic re-engagement of biological processes used during development that enable migrating cells to form tissues. For carcinomas, initiating metastasis is thought to occur through a process called the epithelial-mesenchymal transition (EMT). EMT is the functional consequence of engaging a genetic regulatory network that downregulates the expression of genes associated with an epithelial phenotype and up-regulates genes associated with a mesenchymal phenotype. Breast carcinoma primarily originates from either luminal epithelial cells or basal myoepithelial cells within the mammary gland (10). In contrast to breast cancer, melanoma arises from the oncogenic transformation of melanocytes, which follow a different developmental trajectory along the neural crest than epithelial cells but also involves a process similar to EMT (11). While much of cell specification is imprinted epigenetically via DNA methylation and histone modifications, significant functional changes, such modifications in cell state due to EMT, may occur within these epigenetic constraints. To characterize cell state based on gene expression, supervised methods have been predominantly used for developing gene signatures that characterize the epithelial-mesenchymal transition. While effective, supervised methods can perform poorly if the strategy is based on misinformation, such as sample misclassification or prior biases as to the number of cell states or defining genes. While used less frequently, unsupervised methods for feature extraction and selection are advantageous as they can be data-driven (12). Here, our objective was to develop an unsupervised gene signature capturing this change in phenotype that is tailored to the specific cellular context of breast cancer and melanoma, as an illustrative example.

## Results

### RNA-sequencing provides an estimate of protein abundance

We first asked whether assaying the same genes using different transcriptomics profiling platforms provides the same information. To do this, we compared gene expression levels assayed by either Agilent microarray or by Illumina RNA-sequencing for the same samples (Figure 1). Expression values obtained by RNA-seq are in units of transcripts per million (TPM) while the Agilent Microarray results are in terms of intensity units. Using samples obtained as part of the breast cancer arm of the TCGA, we focused on genes that have been associated with host immunity, as these genes are likely to span a broad dynamic range within these samples. As the TCGA samples are obtained from homogenized bulk samples of tumor and matched normal breast tissue, expression of these genes could be from the malignant cells, like GATA3 expression by breast cancer cells, or from immune cell infiltrates, like the potential expression of IL4 and IL5 by infiltrating T helper type 2 cells.

**Figure 1.**
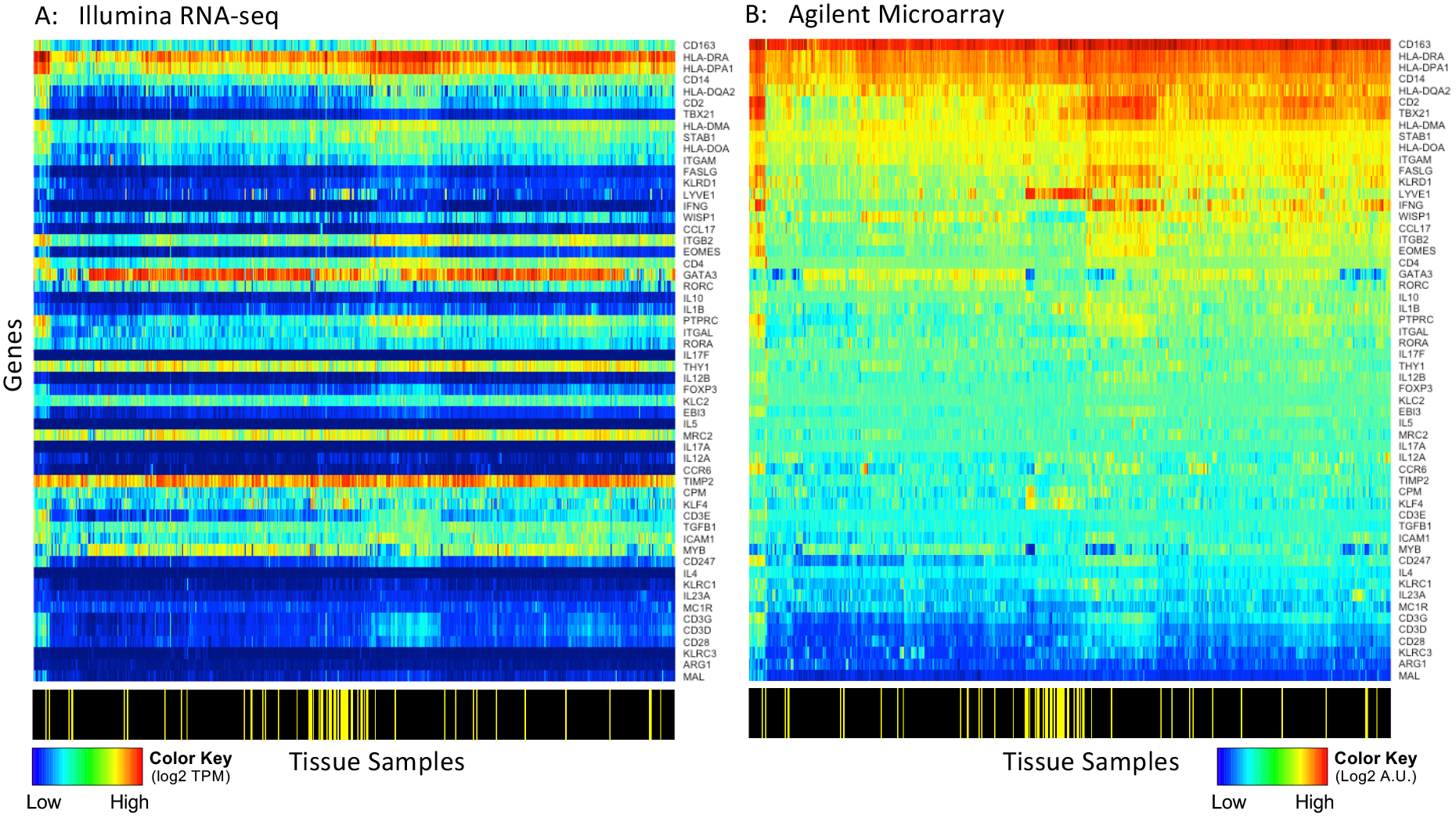
Heatmaps for the expression of a subset of genes in the breast cancer arm of the TCGA study assayed using Illumina RNA-seq (A) and using Agilent microarray (B). Color bar shown at the bottom of the heatmaps indicates samples obtained from tumor tissue (black) versus matched normal tissue (yellow). The genes and samples are similarly ordered in both panels. Values were log2 normalized.

Generally, comparing the same row across the two panels illustrates the poor correspondence between transcript abundance assayed using Agilent microarrays and read counts (TPM) obtained by RNA-seq. A subset of genes, like HLA-DRA and HLA-DPA1, exhibit both high microarray intensity units and read counts while other genes, like TBX21 and FASLG, exhibit high microarray intensity units but have low read counts. In addition, the dynamic range observed among these samples is different depending on the platform used, as illustrated in the heatmap by TBX21 and IL17F. Using Illumina RNA-seq, TBX21 is constrained to the low end of the color spectrum (dark to royal blue) while the dynamic range spans the middle to upper end of the color spectrum (green to red) when assayed using Agilent microarray. Similarly, IL17F transcripts were not detected by RNA-seq in 87% of the samples but the Agilent microarray shows a rather high average intensity with variation among the samples. The difference in average intensities among genes and in variance among samples assayed by these two platforms suggest that the information provided by these two platforms is not entirely the same. The poor correspondence between Agilent two-channel microarray and RNA-seq data has been attributed to differences in ratio (Agilent two-channel microarray) versus non-ratio (RNA-seq) representations of transcript abundance by the platforms (13).

We next asked whether the assayed transcript abundance corresponds to protein abundance. First, we compared RNA-seq counts reported for cell lines associated with the Cancer Cell Line Encyclopedia (CCLE) with protein abundance for the same cell lines measured using Reverse Phase Protein Array (RPPA). We filtered the respective data sets to those cell lines that were reported in both data sets and for genes where there was a positive correlation coefficient greater than 0.36 between read counts in RPKM and normalized RPPA values. From the initial data sets, 283 cell lines and 146 genes were retained for analysis after filtering. Next, we determined whether the pairs of mRNA and protein measurements share a common value for steady-state transcript abundance that corresponds to steady-state protein abundance measured above background. To do this, we applied a protein expression model to each gene measured across the cell lines where protein abundance was assumed to be a saturable function of transcript abundance (Fig. 2A). Using the fitted curve, the threshold of transcript abundance for detecting a change in protein abundance 2.5% above background was back calculated. Example data sets and the corresponding curve fits for the genes CLDN7, AXL, JAG1, and CDH1 are shown in Figure 2B. Interestingly, the peak value in the distribution of calculated threshold values corresponded to 1 RPKM (Fig 2C). We repeated this analysis for transcript abundance assayed by Affymetrix U133+2 microarray, a single-channel approach, using RMA-normalized expression values, where 149 genes were retained for analysis after filtering. Qualitatively, data obtained using the single-channel platform (Affymetrix) exhibited better correspondence with RPPA values than data obtained using Agilent’s two-channel platform. However, the distribution in calculated threshold values were more broadly distributed compared to the RNA-seq results (see red line in Fig 2C, F-test p-value = 0.0045). These results imply that the transcript abundance assayed by RNA-seq provides a higher quality estimate of protein abundance, that is the signal-to-noise is improved, compared with data obtained using a single-channel microarray platform.

**Figure 2.**
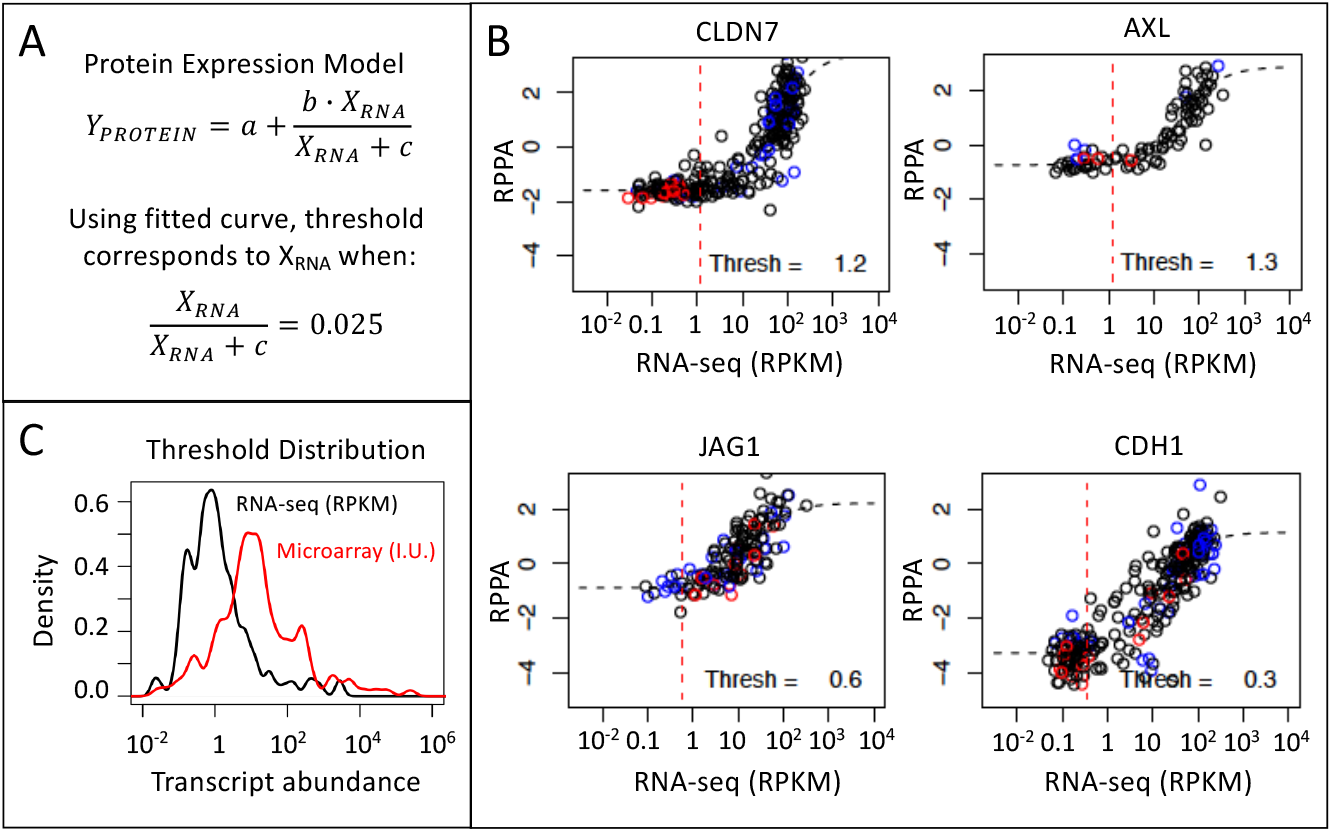
RPPA measurements were used to determine a threshold for biologically significant changes in gene expression. The model for protein dependence on gene expression (A) where representative data (black circles) and model fits (dotted black line) are shown for CLDN7, AXL, JAG1, and CDH1 (B). (C) The distribution in threshold values calculated for all genes assayed by RNA-seq (black curve, n = 146) and by Affymetrix microarray (red curve, n = 149). The reported for transcript abundance units for RNA-seq corresponds to RPKM and intensity units (I.U.) for Affymetrix microarray. In (B), the vertical red dotted line indicates the threshold value and the melanoma and breast cancer cell lines are highlighted by red and blue circles.

Collectively, the common threshold value observed using RNA-seq data has two implications. First, there are some genes that have a high sensitivity of detection using microarrays such that the observed changes may not be functionally important. From Figure 1, it seems that IL17F, TBX21, FASLG, KLRD1, IFNG, CCL17, and IL10 are but a few examples (i.e., high Agilent microarray intensity but very low read counts) in that dataset. Without knowing the detection sensitivity by microarray, traditional approaches using a z-score metric may give equal weight to changes in gene expression driven by a biological signal as to changes dominated by random noise. Second, the threshold value provides a rationale for filtering genes that are likely to have a low in-formation content when developing gene signatures for phenotypes that are not well defined.

### Gene expression patterns in breast cancer cells are captured by a single component

Given the variety of breast cancer subtypes reported in the literature, we next asked how many different genetic regulatory networks (GRN) are at work in breast cancer. GRNs associated with development commonly contain transcription factors that interact via positive feedback such that the target genes are either co-expressed or expressed in a mutually exclusive fashion (14). Given the interest in functional responses, we are focusing on patterns of gene expression in response to signal processing by the genetic regulatory network rather than trying to identify the topology of the GRNs. In motivating this study, we made four assumptions. First, we assumed that oncogenic mutations alter the peripheral control of GRN but do not alter the core network topology, where signals processed by a GRN change cell phenotype by engaging a unique gene expression pattern. Second, malignant cells derived from a particular anatomically-defined cancer represent the diverse ways that hijacking these GRNs can provide a fitness advantage to malignant cells within the tumor microenvironment. Third, culturable tumor cell lines represent a sampling of these ways in which GRNs are hijacked in a particular anatomical location. Fourth, the process of isolating these malignant cells from tumor tissue to generate culturable cell lines does not bias this view. It follows then that the number of different GRNs can be identified by analyzing the transcriptional patterns of genes likely to participate in GRNs among an ensemble of tumor cells lines that share a common tissue of origin. We focused our attention on 780 genes that have been previously associated with the epithelial-mesenchymal transition and related gene sets in MSigDB v4.0. and analyzed the expression of these genes among 57 breast cancer cell lines included in the CCLE database as assayed by RNA-sequencing using a feature extraction/feature selection workflow summarized in Figure 3. To identify coordinately expressed genes, we used Principal Component Analysis (PCA), a linear statistical approach for unsupervised feature extraction and selection that enables the unbiased discovery of clusters of genes that exhibit coherent patterns of expression (i.e., features) that are independent of other gene clusters (15). The relative magnitude of the resulting gene expression patterns can be inferred from the eigenvalues, which is shown in Figure 4. Specifically, PC1 and PC2 captured 65% and 14% of the variance, respectively. Additional principal components each captured less than 4% of the variance.

**Figure 3.**
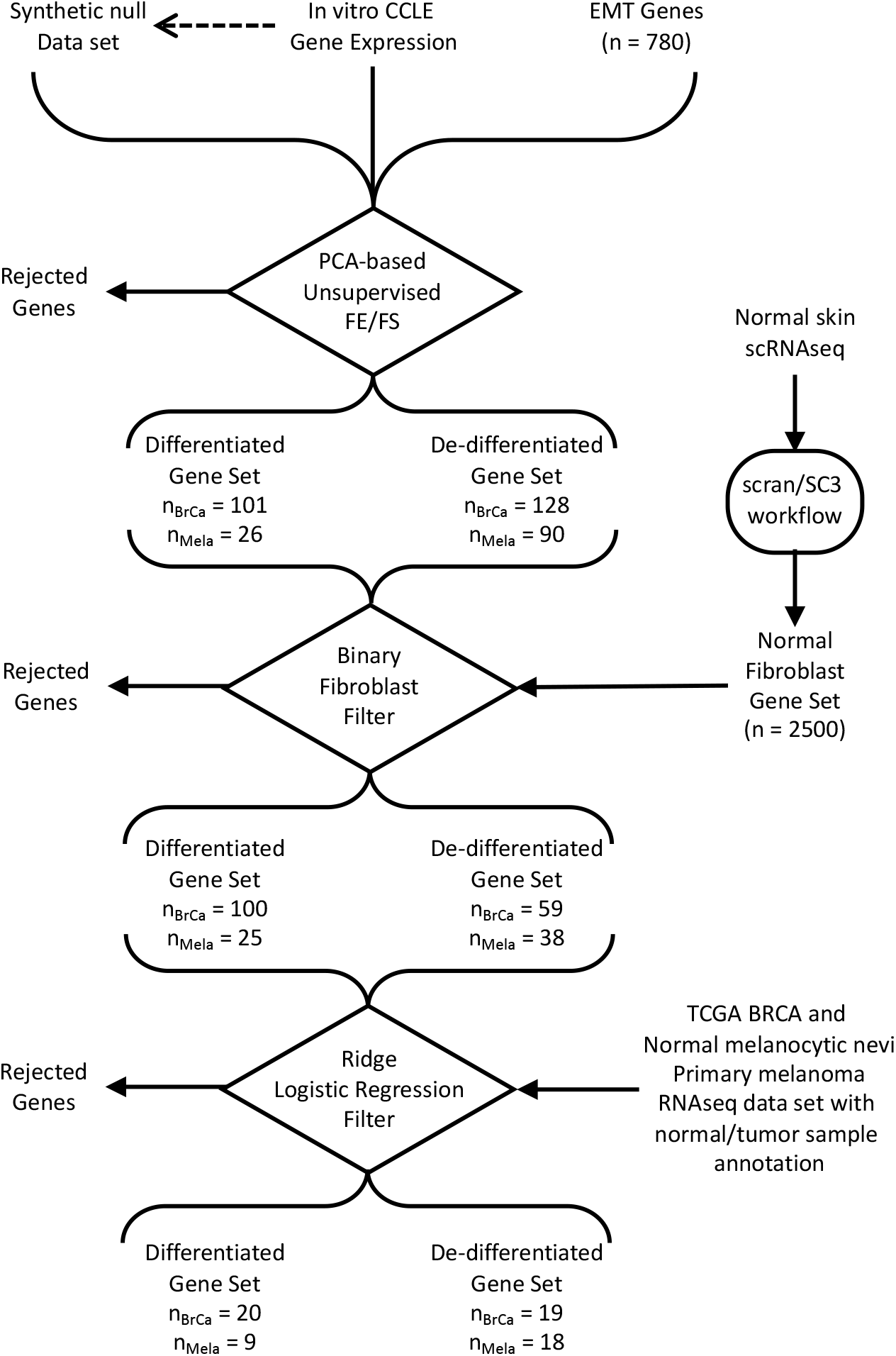
Data workflow for identifying epithelial/differentiated versus mesenchymal/de-differentiated state metrics. Workflow contains three decision points: unsupervised feature extraction (FE)/feature selection (FS) based on PCA, a binary fibroblast filter, and a consistency filter based on Ridge logistic regression of annotated samples.

**Figure 4.**
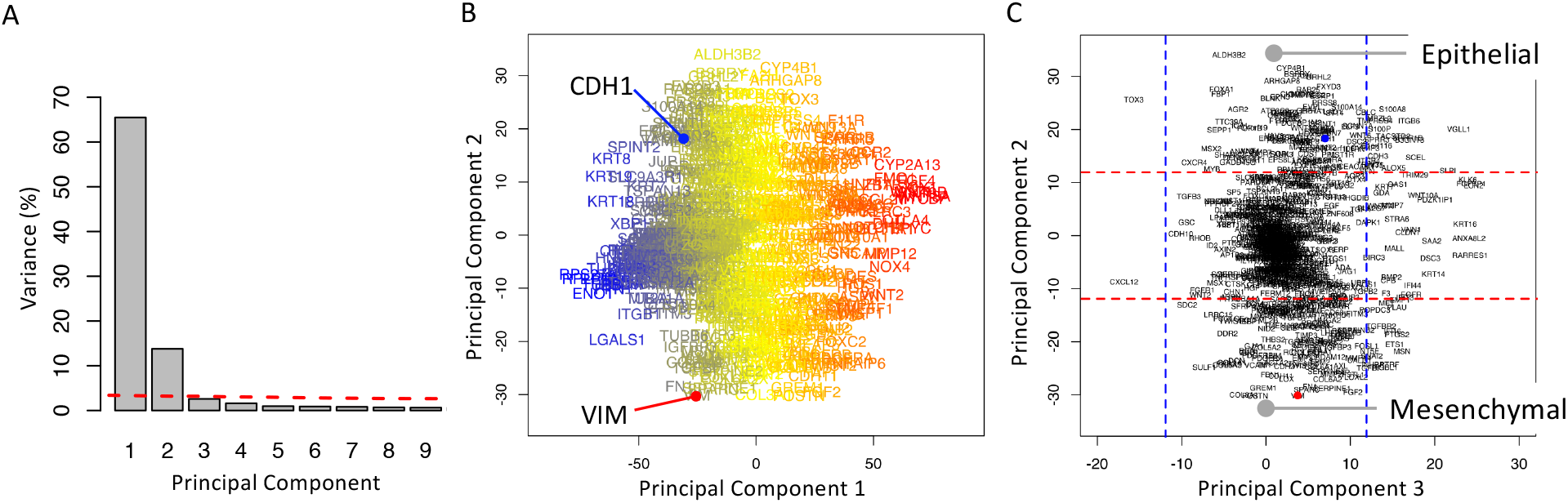
Two opposing gene signatures were identified among the cohort of breast cancer cell lines. (A) Scree plot of the percentage of variance explained by each principal component, where the dotted line corresponds to variance explained by the null principal components. (B) Projection of the genes along PC1 and PC2 axes, where the font color corresponds to the mean read counts among cell lines (blue-yellow-red corresponds to high-medium-low read counts). (C) Projection of the genes along PC2 and PC3 axes, where the dotted lines enclose 95% of the null PCA distribution along the corresponding axis.

One of the challenges with PCA is that no clear rules exist to determine how many principal components to consider, such as a gap statistic in clustering (16). To select an appropriate number of PCs (i.e., features), we established a threshold for determining significance relative to a null distribution. Specifically, we applied the same PCA to a synthetic noise dataset generated from the original data by randomly resampling with replacement the collection of gene expression values and assigning the values to particular gene-cell line combinations. The resulting set of eigenvalues represent the values that could be obtained by random chance if the underlying dataset has no information, which are shown as the dotted red line in Figure 4A. In comparison to the null distribution, only the first two PC were above the threshold. The variance captured by the remaining PCs were below the null PCA distribution suggesting that any potential biological interpretations of these additional PCs could also be explained by random chance. Therefore we focused on the first two PCs.

As variance in read counts is proportional to abundance, gene projections along the PC1 axis were proportional to the average read counts of the corresponding gene among the samples. Measured transcript abundance is proportional to the basal gene expression associated with cell specification and technical artifacts associated with RNA sequencing. Genes were retained for further analysis that were expressed above the 1 RPKM threshold in more that 5% of the cell lines. Next, we focused on the projection of retained genes along principal components 2 and 3. The projections were annotated with horizontal and vertical dotted lines that enclose 95% of the projections from the null distribution. While the majority of the genes were distributed around the origin, a subset of genes were projected along the extreme of the PC2 axis (outside of the dotted horizontal lines) and had no significant projection along the PC3 axis (inside of the dotted vertical lines). The list of genes associated with either the high PC2/null PC3 or the low PC2/null PC3 groups are listed in Supplemental Table S1 and contained 128 and 101 genes, respectively. As the projection of Vimentin (VIM - red dot in Figure 4C) and E-cadherin (CDH1 - blue dot in Figure 4C) were prototypical for these two groups of genes, the high PC2/null PC3 genes were annotated as a mesenchymal signature (i.e., a de-differentiated state) genes and the low PC2/null PC3 group were annotated as an epithelial signature (i.e., a terminally differentiated state). In contrast to supervised approaches that use Vimentin and E-cadherin as the basis to identify associated genes (e.g., (17, 18)), the approach used here is unsupervised whereby the association of Vimentin and E-cadherin with these two opposite groups of genes emerges naturally from the data.

### The Epithelial and Mesenchymal state measures stratify intrinsic subtypes of breast cancer and metastatic potential

Using these two sets of genes, we developed a state metric to quantify the extent of a gene expression signature associated with epithelial differentiation and mesenchymal de-differentiation using a normalized sum over all of the genes associated with a signature. While the PCA results suggest that these two sets of genes are inversely related, the metrics were designed to represent each state independently such that cells that exhibit a pure phenotype would have values of 1 and 0 associated with their respective state metrics and cells with mixed phenotypes could potentially have values of 1 for both state metrics. Next we calculated the state metric values for all of the breast cancer cell lines, where their projections in state space are shown in Figure 5. Interestingly, the breast cancer cell lines largely followed a linear reciprocal relationship between epithelial (E) and mesenchymal (M) states (dotted line in Figure 5) and were segregated by intrinsic PAM50 subtype (19). While HER2, Luminal A, Luminal B, and Basal subtypes all have a high E signature, they progressively increased in their M signature. The Claudin Low subset spanned the greatest range with some expressing a high E and moderate M signatures (e.g., HCC1569, MDAMB361, HMEL) and others with a low E and high M signatures (e.g., BT549 and HS578T). Of note, a subset of the Claudin Low cell lines (e.g., HS742T, HS343T, HS281T, HS606T, and HS274T) with high M and very low E signatures have been suggested by the CCLE to be fibroblastlike (see Cell_lines_annotations_20181226.txt). Functionally, cells with low E and high M signatures had a high propensity for metastasis while the propensity for metastasis was low in cell lines with high E and low M signatures (20).

**Figure 5.**
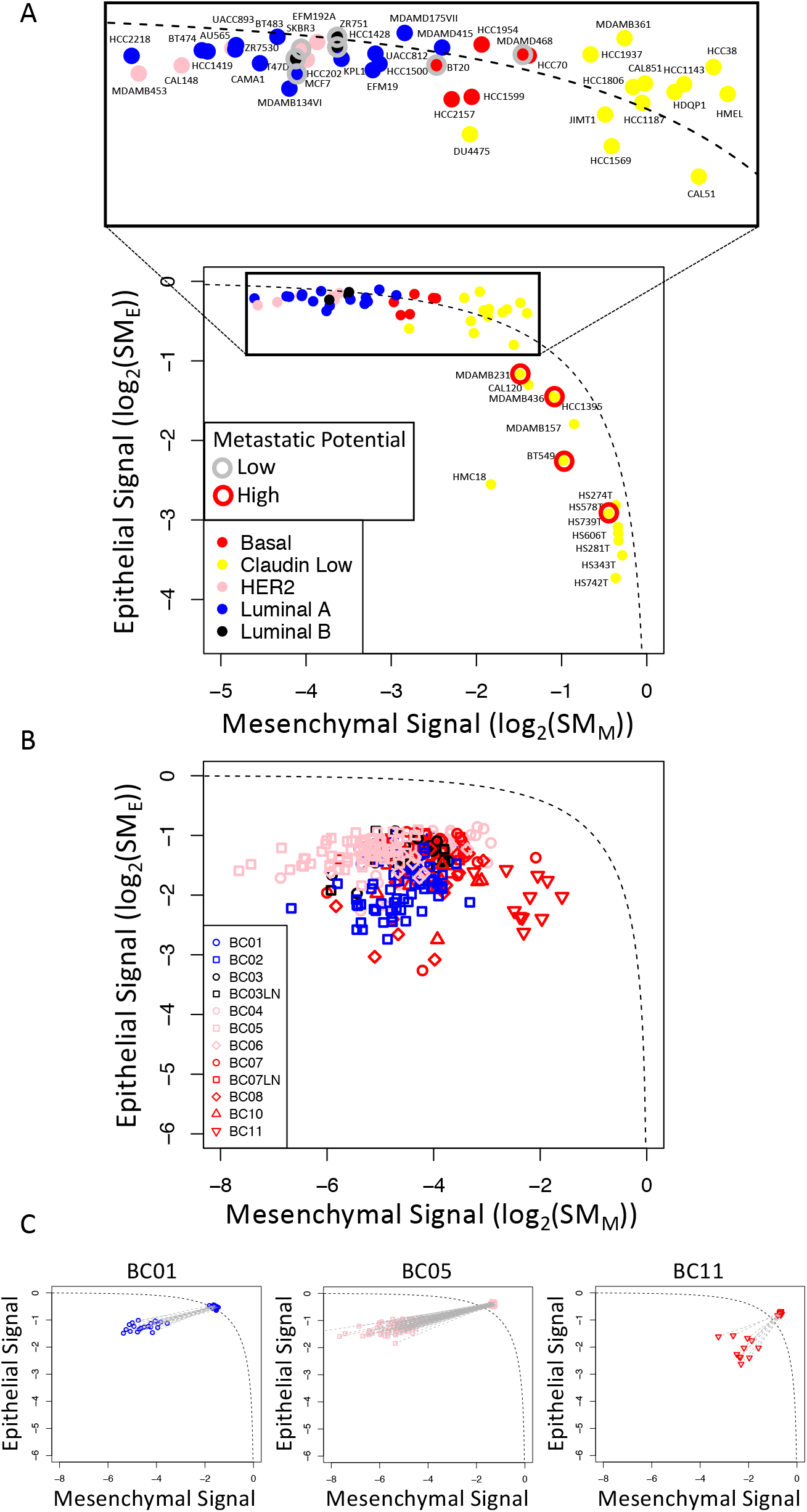
The different subsets of breast cancer were clustered along a reciprocal epithelial to mesenchymal state axes. Log2 projections along the epithelial (*SM*_*E*_) and mesenchymal (*SM*_*M*_) state axes for each breast cancer cell line included in the CCLE (A) and primary breast cancer cells (B and C). Values for *SM*_*E*_ and *SM*_*M*_ were estimated by bulk RNA-seq data for cell lines associated with the CCLE and by scRNA-seq data for primary tumor cells (21). (C) Log2 state projections are compared for primary breast cancer cells as originally reported and with dropout values imputed using the values averaged over the rest of the sample population, where grey lines connect the original state values to state values determine after imputation. Symbols were colored based on previously annotated breast cancer PAM50 subtypes: basal - red, claudin low - yellow, HER2 - pink, luminal A - blue, luminal B - black. In panel A, the metastatic potential of a subset of cell lines were annotated based on a recent study (20): low metastatic potential - grey circle, high metastatic potential - red circle. The dotted line corresponds to a reciprocal relationship between the *SM*_*E*_ and *SM*_*M*_ state metrics (i.e., *SM*_*E*_ = 1 - *SM*_*M*_).

We next assessed the epithelial and mesenchymal state metrics in breast cancer cells assayed using scRNA-seq (21) (see Figure 5B). Similar to the cell lines, the samples were spread across the epithelial to mesenchymal spectrum roughly ordered by their corresponding intrinsic subtype, where HER2 subtype had a high E/low M signature and the basal subtype had the highest M signature without much of a reduction in their E signature. Overall the state values were farther below the reciprocal trendline than any of the cell lines sampled. As gene-level reads by scRNA-seq are frequently missing (i.e., a dropout read)(22), we imputed missing values to assess whether the distribution in the E/M state values were a result of read dropouts (see Figure 5C). While read imputation shifted the cell state metrics toward the reciprocal trendline, the heterogeneity among the cell measurements was lost. Overall, it is unclear whether scRNA-seq measurements can be used to identify biological heterogeneity separately from heterogeneity introduced by technical limits of the assay.

While single-cell methods are rapidly emerging as tool to assay human tissue samples, bulk transcriptomic assays of tumor tissue samples, like those acquired as part of the Cancer Genome Atlas (TCGA) are more abundant. More samples increases the statistical power for identifying clinical, cellular, and genetic correlates of the epithelial-mesenchymal transition. However, applying the epithelial/mesenchymal state metrics to interpret RNA-seq assays of bulk tumor tissue samples requires some additional filtering steps as bulk RNA-seq measurements averages over the heterogeneous normal and malignant cell types present within the tissue. In terms of a gene signature for the epithelial-mesenchymal transition, many of the genes commonly associated with acquiring mesenchymal function are also associated with fibroblasts, a relatively common cell type in epithelial tissues. Thus, an enrichment of genes associated with the epithelial-mesenchymal transition may be explained solely by a shift in the prevalence of fibroblasts within the tissue sample. To deconvolute fibroblast genes from the state metrics, we obtained a list of 2500 genes that were uniquely associated (Area under ROC curve *>* 0.5) with a cluster annotated as fibroblasts using scRNA-seq data obtained from a digested normal skin sample obtained from a human female. This cluster contained about 1/3 of the cell samples measured within the CD45-negative population of the digested skin sample (see Supplemental Figure S1). Using this fibroblast gene list, overlapping genes were removed from the state metrics and highlighted in yellow in Supplemental Table S1. All but one of the genes removed were contained within the mesenchymal gene list.

Gene expression assayed from a bulk tissue sample reflects the combined contributions of non-malignant cells plus the changes induced by oncogenic transformation, and reciprocal changes due to de-differentiation among malignant cells. Observable changes depend on the relative contributions of each cell source. As the unsupervised PCA analysis of the cell line data suggested that genes associated with EMT can be revealed by identifying a reciprocal pattern of gene expression, we performed Ridge logistic regression using the sample annotation to obtain regression coefficients for the list of EMT genes that passed the fibroblast filter (n = 158). The regression coefficients were used to filter the list of EMT genes for consistency with the reciprocal gene signature identified in the CCLE analysis. Genes that passed the consistency filter were used to define the epithelial and mesenchymal state metrics for bulk tissue samples. Of note, E-cadherin (CDH1) and CEACAM1 were associated with the epithelial state metric while N-cadherin (CDH2), Wnt-inducible signaling pathway protein 1 (WISP1/CCN4), and matrix metallopeptidase 3 (MMP3) were associated with the mesenchymal state metric. The list of genes associated with the corresponding state metrics are summarized in Supplemental Table S2.

Next, we projected the tissue samples obtained as part of the breast cancer arm of the TCGA in EMT space using the two tissue-based state metrics. Similar to the CCLE analysis, all samples clustered along the reciprocal *SM*_*E*_ versus *SM*_*M*_ line but exhibited greater dispersion. Samples obtained from normal breast tissue clustered separately from breast cancer samples (Figure 6), with normal breast tissue samples having the highest values for the epithelial state and lower values, on average, for the mesenchymal state. Among the different clinical breast cancer subtypes, the median value for *SM*_*E*_ progressively decreased from ER/PR+ (luminal), HER2+, and triple negative (TN: ER-/PR-/HER2-) subtypes. The mesenchymal projections were about equal for both ER/PR+ and HER2+ subsets and higher than the TN samples. We note that, while the HER2+ samples were roughly equal to the ER/PR+ along the *SM*_*M*_ axis, the HER2+ samples were lower than the ER/PR+ samples along the *SM*_*E*_ axis, which aligns with clinical observations. For instance, patients with HER2+ subtype of breast cancer are at increased risk for developing metastatic lesions compared to TN and luminal sub-types (23). The two different state metrics seem to capture gene signatures that helps anchor a cell to its’ designated location within the tissue and that promotes active migration, respectively. In other words, reducing *SM*_*E*_ corresponds to raising the anchor and increasing *SM*_*M*_ corresponds to hoist-ing the sail. In summary, both cell-level and tissue-level EMT state metrics provide an estimate of metastatic potential and a digital measure of malignant cell differentiation state in the context of breast cancer.

**Figure 6.**
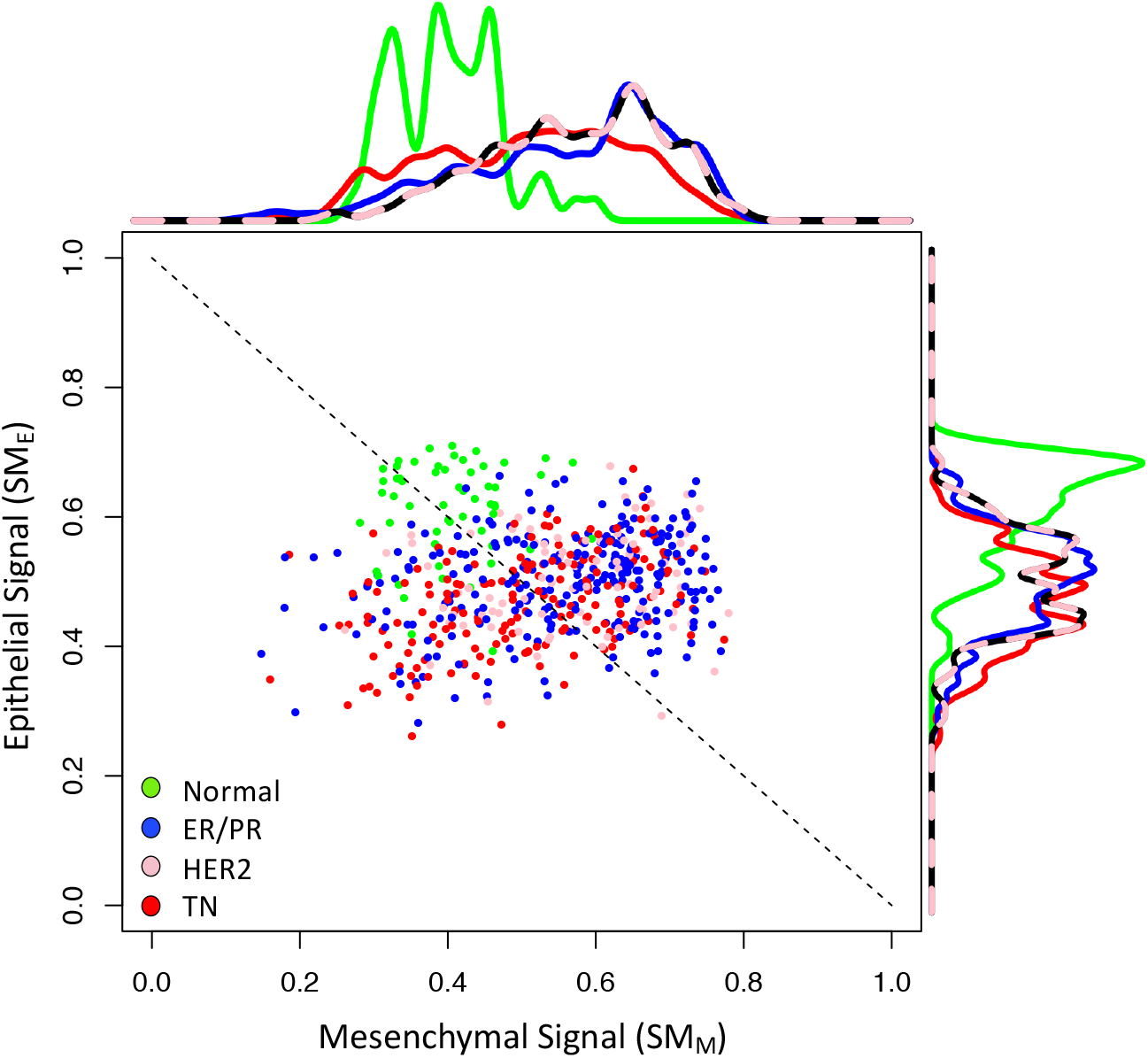
The samples from normal breast tissue and breast cancer were clustered separately along a reciprocal epithelial to mesenchymal state axes. Using EMT genes that passed the gene filter workflow, each sample contained within the breast cancer (BrCa) arm of the TCGA was projected along the epithelial (*SM*_*E*_) versus mesenchymal (*SM*_*M*_) state axes using the corresponding bulk RNA-seq data. Symbols were colored based on normal breast tissue (green) or clinical breast cancer subtype: ER/PR+ - blue, HER2 - pink, triple negative (TN) - red. The dotted line corresponds to a reciprocal relationship between the *SM*_*E*_ and *SM*_*M*_ state metrics (i.e., *SM*_*E*_ = 1 - *SM*_*M*_).

### Gene expression patterns in melanoma cells are also captured by a single component

Using the same feature extraction/feature selection workflow as the breast cancer analysis (Figure 3), we applied principal component analysis to the expression of EMT-related genes assayed in an ensemble of 56 melanoma cell lines associated with the Cancer Cell Line Encyclopedia (Figure 7). We focused on the first two principal components, PC1 and PC2, that captured 80% and 6% of the variance, respectively. Additional principal components each captured less than 4% of the variance. PC1 captured the variance associated with read abundance, as gene projections along the PC1 axis were proportional to the average read counts among the samples. Vimentin (VIM) and fibronectin (FN1) were two of the most highly expressed genes while members of the Wnt family were some of the genes with low expression (e.g., WNT1, WNT6, WNT8B, WNT3A, WNT8A, WNT9B). Genes retained for further analysis were expressed above the 1 RPKM threshold in more than 5% of the cell lines.

**Figure 7.**
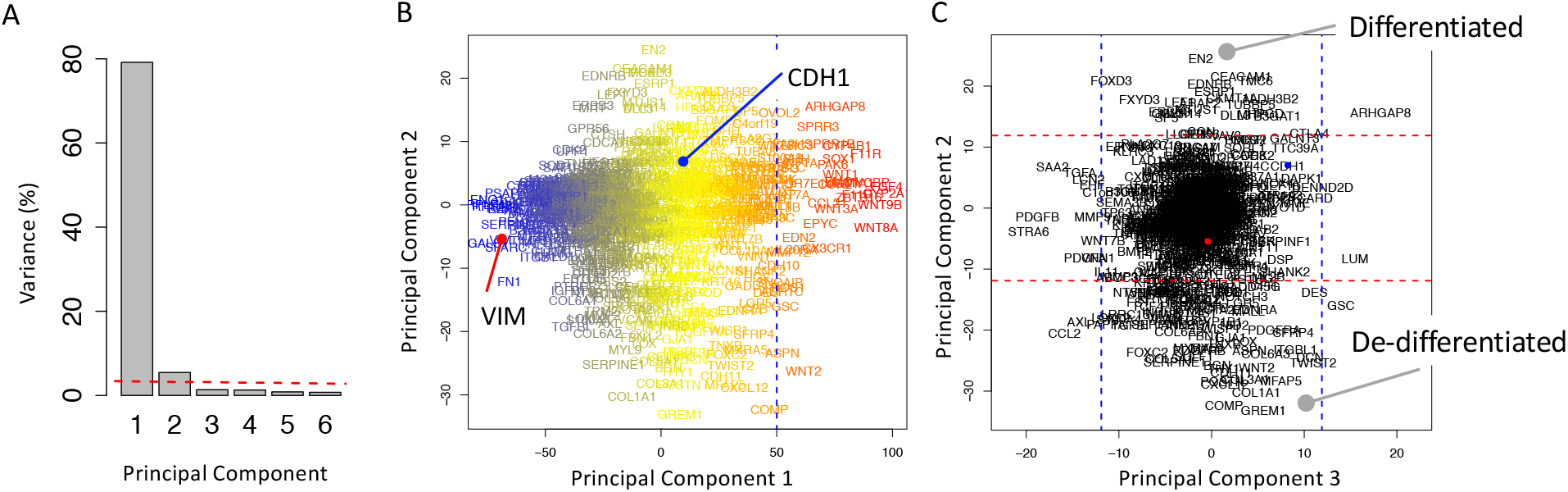
Two opposing gene signatures were identified among the cohort of melanoma cell lines. (A) Scree plot of the percentage of variance explained by each principal component, where the dotted line corresponds to variance explained by the null principal components. (B) Projection of the genes along PC1 and PC2 axes, where the font color corresponds to the mean read counts among cell lines (blue-yellow-red corresponds to high-medium-low read counts). (C) Projection of the genes along PC2 and PC3 axes, where the dotted lines enclose 95% of the null PCA distribution along the corresponding axis.

Next, we focused on the projection of retained genes along PC2 and PC3 axes. The projections were annotated with horizontal and vertical dotted lines that enclose 95% of the projections from the null distribution. While the majority of the genes were distributed around the origin, a subset of genes were projected along the extreme of the PC2 axis (outside of the dotted horizontal lines) and had no significant projection along the PC3 axis (inside of the dotted vertical lines). The list of genes associated with either the high PC2/null PC3 or the low PC2/null PC3 groups are listed in Supplemental Table S1 and contained 26 and 90 genes, respectively. In contrast to the breast cancer results, the projection of Vimentin (VIM - red dot in Figure 7C) and E-cadherin (CDH1 - blue dot in Figure 7C) were not associated with either of these two groups of genes. As the high PC2/null PC3 group included MITF, a master regulator of melanocyte differentiation, and the low PC2/null PC3 group included a number of EMT-related genes (e.g., FN1, TCF4, ZEB1, TWIST2, and WISP1), these two gene sets were annotated as a terminally differentiated signature (i.e., an epithelial-like state) and a de-differentiated signature (i.e., a mesenchymal-like state), respectively.

Projections of the melanoma cell lines in differentiation state space were calculated using the two state metrics (Figure 8). Similar to the breast cancer cell lines, the melanoma cell lines largely followed a linear reciprocal relationship between terminally differentiated (*SM*_*T*_) and de-differentiated (*SM*_*D*_) states (dotted line in Figure 8). The majority of cell lines exhibited primarily a terminally differentiated signature with some expression of de-differentiated genes while only a small subset of the cell lines exhibited primarily a de-differentiated signature. The gene signatures for single melanoma cells were also highly heterogeneous due to dropout of gene reads.

**Figure 8.**
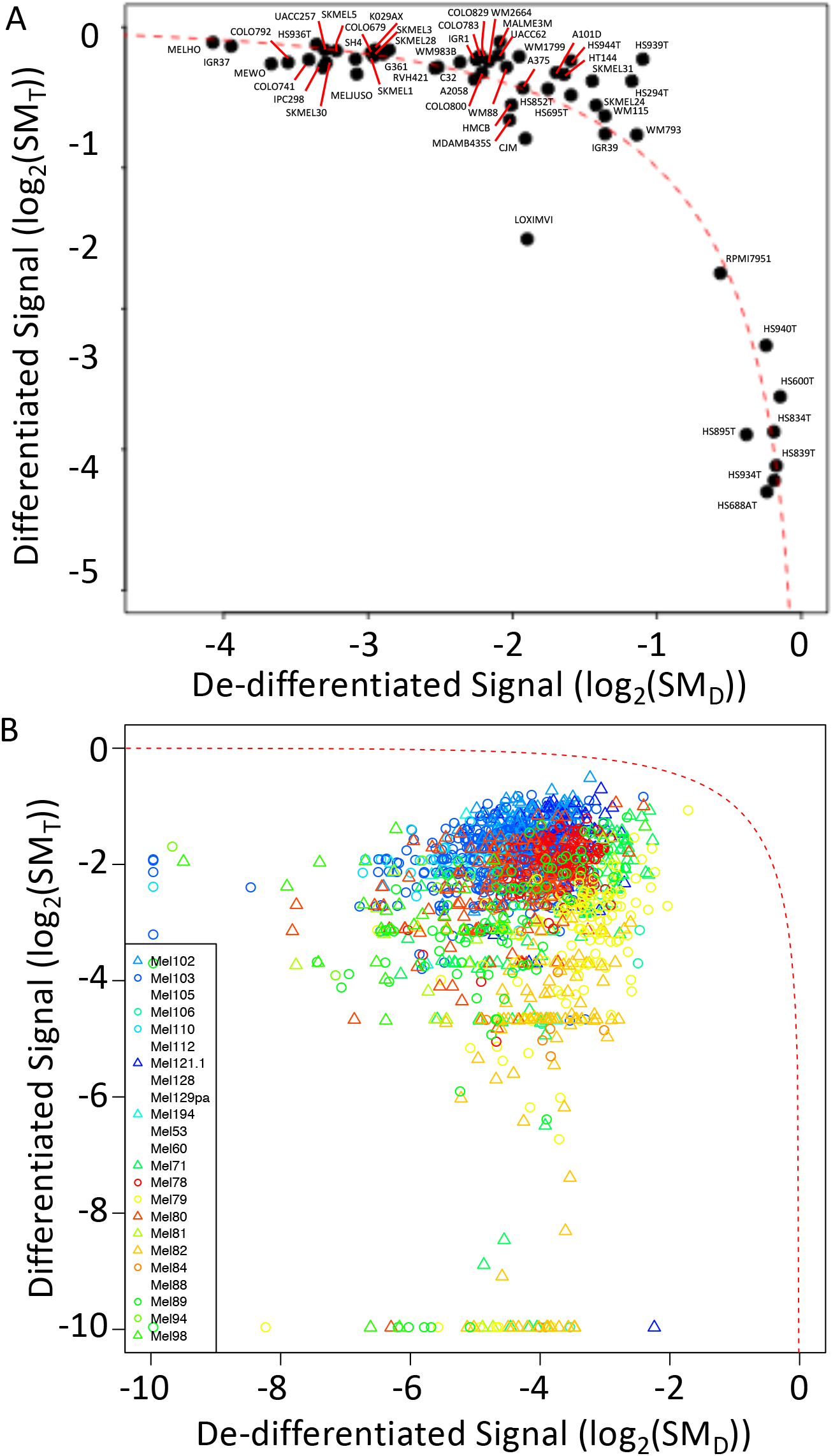
Melanoma cell lines and primary single melanoma cells are distributed along path between extremes in differentiation states. Projections along the terminally differentiated (*SM*_*T*_) versus de-differentiated (*SM*_*D*_) state axes for each melanoma cell line included in the CCLE (A) and primary melanoma cells (B). Values for the terminally differentiated and de-differentiated state metrics were estimated by RNA-seq data for cell lines associated with the CCLE and by scRNA-seq data for primary melanoma cells. Symbols for primary melanoma cells were colored differently for each patient sample. The dotted line corresponds to a reciprocal relationship between the *SM*_*T*_ and *SM*_*D*_ state metrics (i.e., *SM*_*T*_ = 1 - *SM*_*D*_).

Using state metrics refined for use with tissue samples (see Supplemental Table S2), samples acquired from benign melanocytic nevi and untreated primary melanoma tissue were projected onto the state space. Of note, CEACAM1 and MITF were associated with the differentiated state and three genes - CEACAM1, CGN, and HPGD - were shared with the breast cancer epithelial state metric. The de-differentiated state metric had five genes - WISP1/CCN4, EDNRA, FOXC2, SERPINE1, and SPOCK1 - that were shared with the breast cancer mesenchymal state metric. While samples were more narrowly distributed in state space compared to the cell lines (Figure 9), all of the benign nevi exhibited higher terminally differentiated (*SM*_*T*_) and tended to have lower de-differentiated values (*SM*_*D*_). The samples from primary melanoma were color-coded based on the annotated Breslow’s depth, where higher values were associated with lower terminal differentiation scores. Using Breslow’s depth as a surrogate measure of metastatic potential (24), tissue-level EMT state metrics provide an estimate of metastatic potential and a digital measure of malignant cell differentiation state in the context of melanoma.

**Figure 9.**
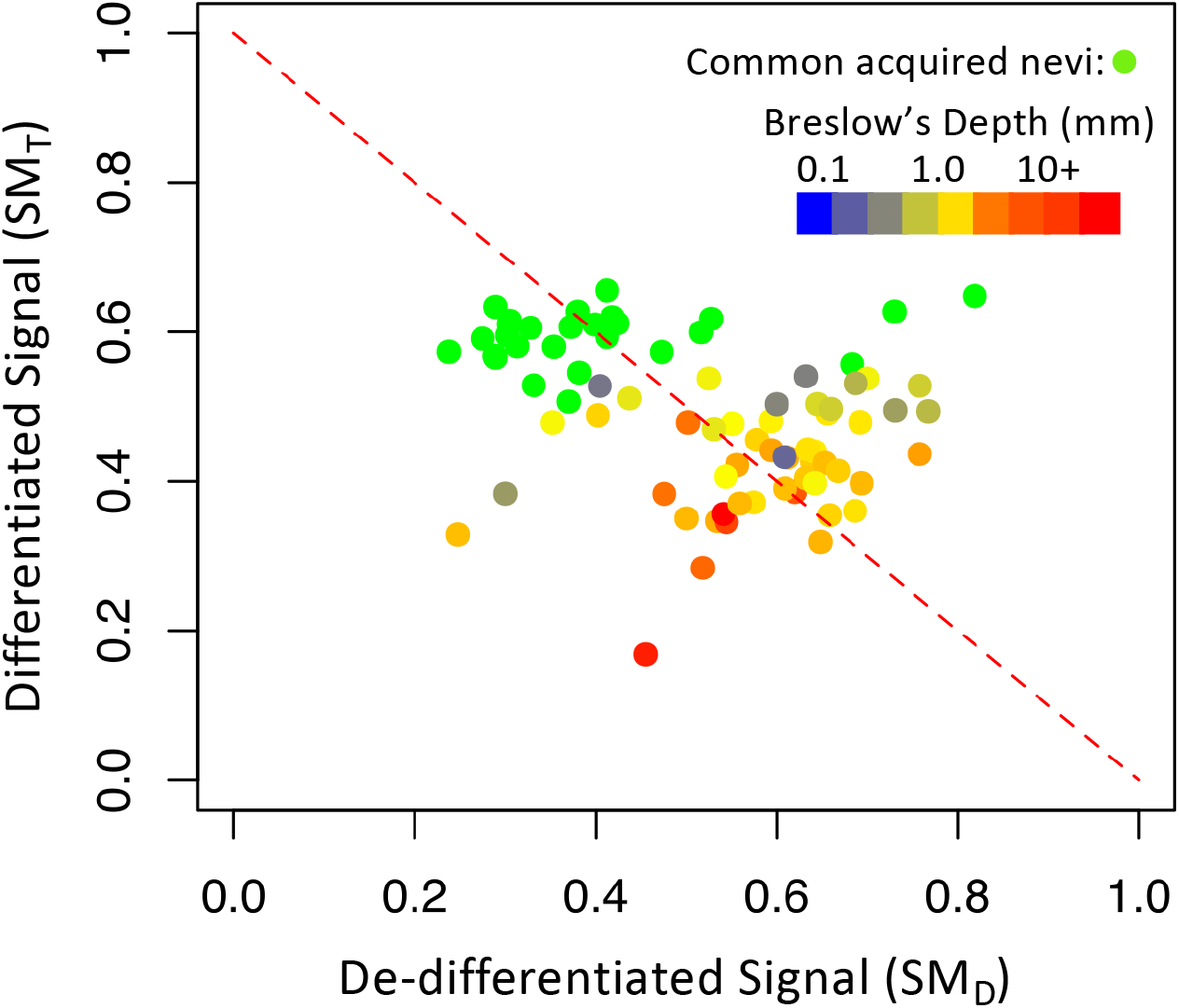
Gene expression patterns associated with benign melanocytic nevi and primary melanoma tissue samples are distributed along path between extremes in differentiation states. Projections along the terminally differentiated (*SM*_*T*_) versus de-differentiated (*SM*_*D*_) state axes for 78 tissue samples obtained from common acquired melanocytic nevi (n = 27, green circles) and primary melanoma (n = 51). The primary melanoma samples are colored based on the Breslow’s depth (blue: 0.1 mm to red: 10+ mm). The dotted line corresponds to a reciprocal relationship between the *SM*_*T*_ and *SM*_*D*_ state metrics (i.e., *SM*_*T*_ = 1 - *SM*_*D*_).

**Figure 10.**
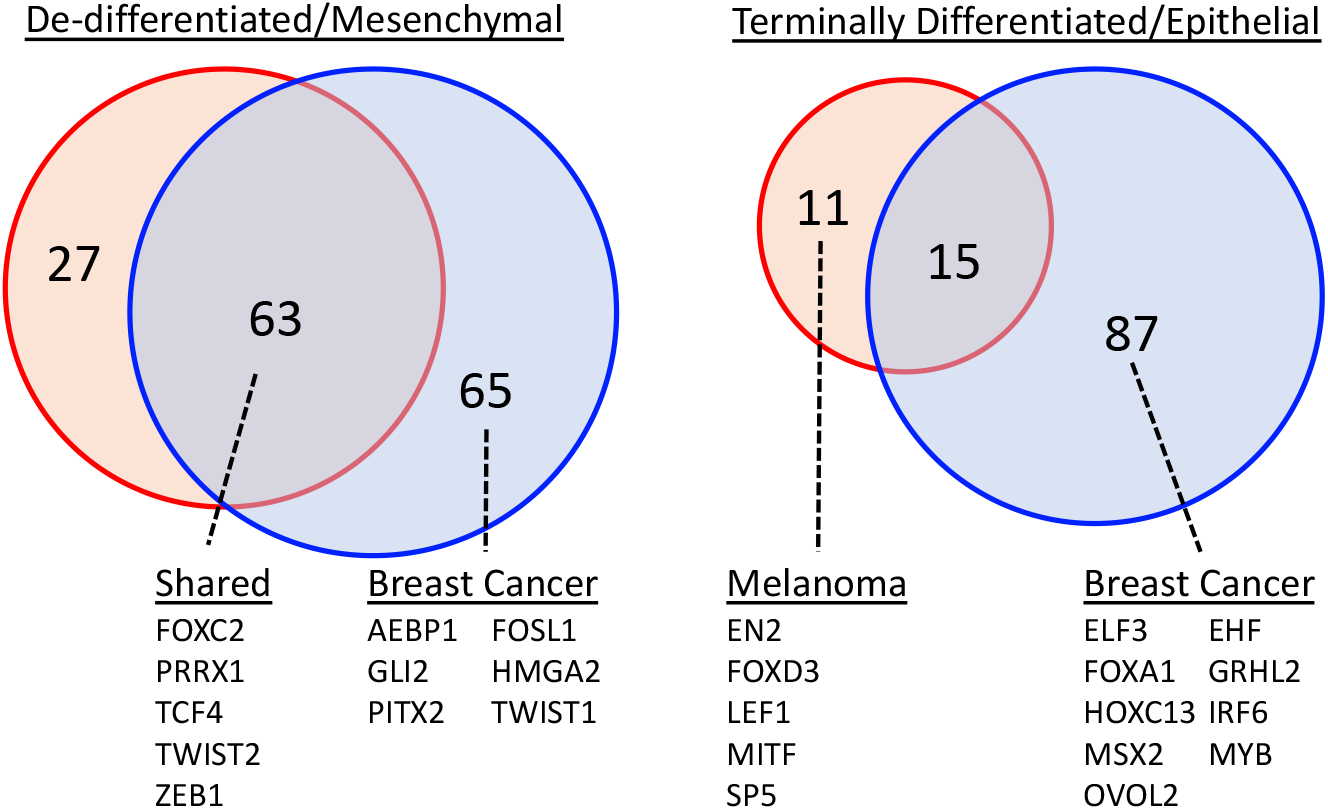
Venn diagram illustrating overlap in genes contained in the opposing state metrics for terminally differentiated/epithelial versus de-differentiated/mesenchymal extracted from breast cancer (blue circle) and melanoma (red circle) cell lines. The subset of the genes listed below the Venn diagram were annotated with transcription factor GO terms.

### Terminal differentiation is associated with distinct gene signatures while de-differentiation seems to engage common gene regulatory networks

The separate gene signatures generated for breast cancer cells and melanoma cells using an unsupervised approach provide an opportunity to identify unique and shared aspects of the genetic regulatory mechanisms underpinning cell specification, as summarized in Figure 9. Specifically, we used DAVID to identify genes with transcription factor activity using the GOTERM_MF_ALL: Sequence_specific_DNA_binding + UP_Keywords:DNA_binding. In the breast cancer cell lines, nine transcription factors were upregulated in cells with a terminally differentiated phenotype, including GRHL2 and OVOL2 that have been associated with enforcing epithelial differentiation (25). Correspondingly, five transcription factors were upregulated in melanoma cells, including MITF that is essential for melanocyte differentiation (26). Interestingly, there was no overlap in the genes with transcription factor activity in the two differentiated cell signatures. In contrast, melanoma and breast cancer cell lines that exhibited a de-differentiated phenotype shared five transcription factors, including TWIST2 and ZEB1. De-differentiation in breast cancer cell lines were also associated with an additional six transcription factors, including TWIST1 (27). Overall, the analysis of these transcription factors is consistent with specificity in phenotype as a consequence of engaging gene regulatory networks unique to a specialized cell subset while de-differentiation seemed to engage common gene regulatory networks that facilitate the loss of cell specificity.

## Discussion

Here we used an unsupervised feature extraction and selection approach based on principal component analysis and resampling to identify state metrics for the epithelial-mesenchymal transition in breast cancer and melanoma. Given the importance for identifying patients with tumors likely to metastasize, a number of gene signatures have been developed to predict the prevalence of tumor cells with a epithelial-mesenchymal transition signature (17, 18, 28–30). Supervised approaches are most common (17, 18, 28, 29), where samples are classified a priori. For instance, Koplev et al. (29) develop gene signatures that average over all anatomical locations while Levine and coworkers (28, 31) classify training samples a priori into one of three cell states: epithelial, mesenchymal, or hybrid E/M. Rokavec et al. generate features based on co-expression with E-cadherin and Vimentin (18). While effective, supervised methods can perform poorly if the strategy is based on misinformation, such as sample misclassification or prior biases as to the number of cell states or defining genes. We also note that state metrics developed using microarray technology (e.g., (17, 29)) are not likely relevant for interpreting data based on RNA sequencing, given the unclear relation between transcriptome and protein abundance as assayed using microarray technology. While rarely used, the data-driven nature of unsupervised methods for feature extraction and selection are attractive (12). For instance, Umeyama et al. used an unsupervised approach for feature extraction to identify genes associated with metastasis (32). To illustrate this data-driven approach, we have focused on breast cancer and melanoma, where metastatic dissemination to vital organs are key limiters of patient survival. In summary, we hope that our developed state metrics find use alongside other digital cytometry tools to better understand how oncogenic transformation alters the immune contexture within the tumor microenvironment.

## Methods

### ’Omics Data

Transcriptomics profiling of the same samples using both Agilent microarray and Illumina RNA sequencing for the breast cancer arm (BRCA) of the Cancer Genome Atlas was downloaded from TCGA data commons. Values for gene expression, expressed in RPKM for RNA-seq and gene-centric RMA-normalized data for Affymetrix U133+2 microarray, for the cell lines contained within the Cancer Cell Line Encyclopedia were downloaded from the Broad data commons (Website: https://portals.broadinstitute.org/ccle Files: CCLE_RNAseq_081117.rpkm.gct accessed 12/22/2017 and CCLE_Expression_Entrez_2012-10-18.res accessed 6/15/2018). Reverse phase protein array (RPPA) results for the cancer cell lines were obtained from the M.D. Anderson proteomics website (Website: https://tcpaportal.org/mclp/ File: MCLP-v1.1-Level4.txt accessed 6/15/2018) (6). Single-cell gene expression (scRNA-seq) for breast cancer and melanoma cells expressed in TPM were downloaded from the Gene Expression Omnibus (GEO) entries GSE75688 and GSE72056, respectively. 10X Genomics scRNA-seq data for CD45-negative cells digested from a normal human female skin sample and expressed in counts of gene-level features was downloaded from European Bioinformatics Institute (EMBL-EBI) ArrayExpress entry E-MTAB-6831. RNA-seq data expressed in counts assayed in samples acquired from benign melanocytic nevi and untreated primary melanoma tissue and associated annotation were downloaded from GEO entry GSE98394.

### Non-linear regression of protein abundance to mRNA expression

All data was analyzed in R (V3.5.1) using the ‘stats’ package (V3.5.1). For each gene where complementary CCLE transcriptomic and RPPA data exist and for which their correlation coefficient was above 0.36, the non-linear function,

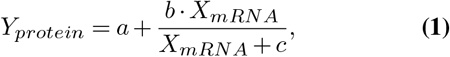

was regressed using the *nls* function to the resulting protein (*Y_protein_*) and transcript (*X_mRNA_*) abundance data. As the RPPA values are normalized, the parameters *a* and *b* represent the background value and maximum detectable increase above background, respectively, while the parameter *c* represents the midpoint in transcript abundance within the dynamic range of the assay. A minimum in the summed squared errors between model-predicted and observed RPPA values were used to determine the optimal values of the model parameters. Using the optimal values, a threshold was estimated independently for each gene based on the transcript abundance that yields a 2.5% increase in protein abundance above background. The regression was repeated using both RNA-seq and Affymetrix transcriptomics data.

### Statistical analysis for cell-level signatures

Principal component analysis (PCA) was performed on log base 2 transformed RPKM values using the *prcomp* function on the CCLE RNA-seq data, which was filtered to 780 genes previously associated with epithelial-mesenchymal transition. The collective list of genes were assembled from prior studies (17, 33–37) and additional gene sets from MSigDB V4.0 including: “EPITHELIAL TO MESENCHYMAL TRANSITION” and “REACTOME TGF BETA RECEPTOR SIGNALING IN EMT EPITHELIAL TO MESENCHYMAL TRANSITION". PCA was applied to the genes to extract the features, where the resulting eigenvectors capture the relative influence of a gene’s expression on a specific principal component and the eigenvalues represent how much information contained within the dataset is captured by a specific principal component. Drawing upon conventional hypothesis testing where significance is established by rejecting the null hypothesis that experimental observations could be explained by random chance, we used a resampling approach to establish a null hypothesis related to the eigenvalues, that is to determine the true rank of the noisy expression matrix. The resampling approach involved repetitively applying PCA (n = 1000) to a synthetic noise dataset with the same dimensions that was generated from the original data by randomly resampling with replacement from the collection of gene expression values and assigning the values to particular gene-cell line combinations. The resulting distribution of eigenvalues and eigenvectors represent the values that could be obtained by random chance if the underlying dataset has no information (i.e., the null PCA distribution). Principal components with eigenvalues greater than the null PCA distribution were used to define the principal subspace for subsequent analysis, that is the selection of features. Similarly, the distribution in the projection of genes within the null PCA space were used to determine whether the projection of a gene along a particular PC axis was explained by random chance or not by setting thresholds along the PC2 and PC3 axes that enclosed 95% of the null PCA space. The PC projection of genes relative to the null PCA space was used to refine the extracted features. A metric was developed to estimate the extent that a cell exhibits a gene signature corresponding to a “Epithelial/Terminally Differentiated" versus “Mesenchymal/Dedifferentiated" state. The state metrics (*SM*) quantify the cellular state by averaging over a normalized expression level of each gene in the signature (*reads*_*i*_, expressed in TPM) according to the formula:

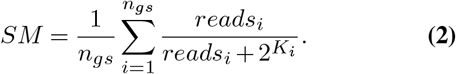

The genes included in a signature with their corresponding *K*_*i*_ values are listed in Supplemental Table S1 and *n*_*gs*_ corresponds to the number of genes within a signature. The *K*_*i*_ values were estimated by clustering the log2 expression of each gene into two groups using the k-means method and the value was set as the mid-point in expression between the two groups.

### Statistical analysis for tissue-level signatures

Genes differentially expressed in normal epithelial fibroblasts were obtained by analyzing single-cell RNA-seq data of normal skin obtained using a Genomics 10x platform and a bioinformatics workflow based on the scater (V1.12.2) and SC3 (V1.12.0) packages in R. Briefly, scRNA-seq data were filtered to retain samples that had less than 50% of the reads in the top 50 genes and to remove outlier samples based on PCA analysis. Gene-level features were limited to those that were expressed at greater than 1 count in more than 10 cell samples. Read depth was normalized using a variant of CPM contained within the *scran* (V1.12.1) package, which develops a sample-specific normalization factor repetitive sample pooling followed by deconvoluting a sample-specific factor by linear algebra. Following from Davidson et al. (38), fibroblasts were annotated based on co-expression of COL1A1 and COL1A2. Samples were clustered and genes differentially associated with each cluster were identified using the *SC3* workflow (V1.14.0) using default parameters (see Figure S1).

Prior to logistic regression analysis, TCGA BRCA data and the benign nevi and melanoma data were filtered to remove sample outliers and normalized based on housekeeping gene expression (39). Using normal versus tumor annotation associated with the data, ridge logistic regression was performed on log base 2 transformed TPM and median-centered values using the *glmnet* package (V2.0-18), which was limited to EMT-related genes identified in the CCLE analysis and not associated with normal fibroblasts. To minimize overfitting, ridge logistic regression was repeated 500 times using a subsample of the original data set using the genes associated with each signature separately. In each iteration, the samples were randomly assigned in an 80:20 ratio between training and testing samples. Regression coefficients were captured for each iteration using a lambda value that minimized the misclassification error of a binomial prediction model estimated by cross-validation. Accuracy was assessed using the testing samples. Genes were determined to have a consistent expression pattern if greater than 95% of the distribution in regression coefficients had the correct sign. Similarly to the cell-level analysis, state metrics were developed for bulk tissue-level RNA-seq measurements to estimate the extent that a tissue sample exhibits a gene signature corresponding to a “Epithelial/Terminally Differentiated” versus “Mesenchymal/De-differentiated” state. The genes included in a signature and their corresponding *K*_*i*_ values are listed in Supplemental Table S2.

## ACKNOWLEDGEMENTS

This work was supported by National Science Foundation (NSF CBET-1644932 to DJK) and National Cancer Institute (NCI 1R01CA193473 to DJK). The content is solely the responsibility of the authors and does not necessarily represent the official views of the NSF or NCI.

## AUTHOR CONTRIBUTIONS

These contributions follow the International Committee of Medical Journal Editors guidelines: http://www.icmje.org/recommendations/. Conceptualization: DJK; Study Design: DJK; AT;; Data Analysis: DJK; AT; Data Interpretation: DJK; Funding acquisition: DJK; Methodology: DJK; AT; Project administration: DJK; Software: DJK; AT; Supervision: DJK; Writing – original draft: DJK; Writing – review & editing: all authors.

---

**Table S1.**
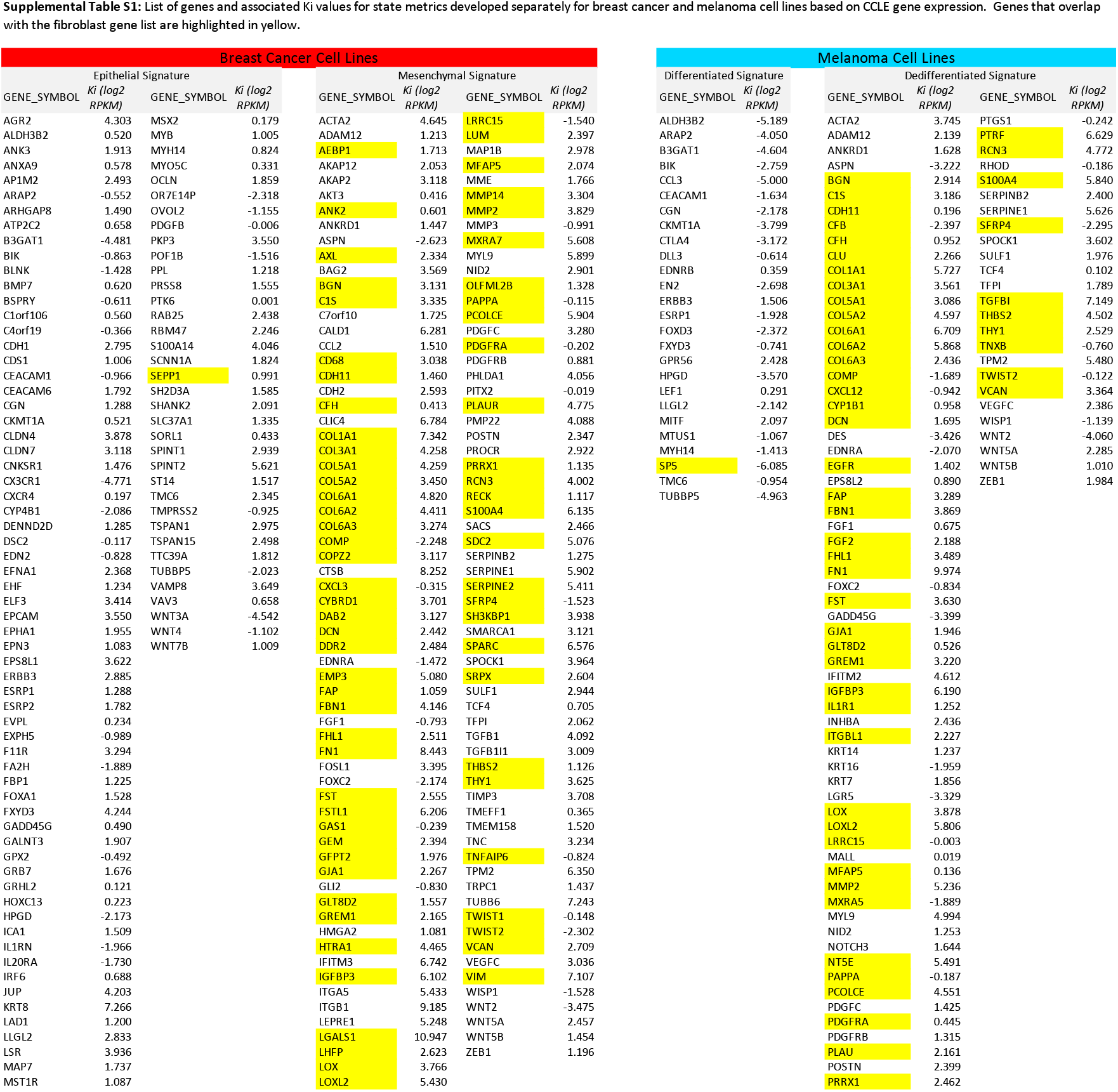
List of genes and corresponding *K_i_* values for state metrics developed separately for breast cancer and melanoma cell lines based on CCLE gene expression. Genes that overlap with the fibroblast gene list are highlighted in yellow.

**Table S2.** List of genes and associated Ki values for refined state metrics based on TCGA breast cancer tissue samples and tissue samples of common acquired melanocytic nevi and primary melanoma. Genes that overlap in the state metrics between breast cancer and melanoma are highlighted in green.

| TCGA Breast Cancer Tissue Samples |  |  |  | Melanocytic Nevi and Melanoma Tissue Samples |  |  |  |
| --- | --- | --- | --- | --- | --- | --- | --- |
| Epithelial Signature |  | Mesenchymal Signature |  | Differentiated Signature |  | De-differentiated Signature |  |
| GENE_SYMBOL | Ki (log2 TPM) | GENE_SYMBOL | Ki (log2 TPM) | GENE_SYMBOL | Ki (log2 TPM) | GENE_SYMBOL | Ki (log2 TPM) |
| ALDH3B2 | 6.539 | ADAM12 | 5.020 | ARAP2 | 7.068 | ACTA2 | 5.441 |
| ANK3 | 3.420 | ASPN | 6.415 | CEACAM1 | 3.160 | DES | 0.960 |
| B3GAT1 | -1.558 | CDH2 | 1.919 | CGN | 5.204 | EDNRA | 4.051 |
| BMP7 | 2.158 | CLIC4 | 8.028 | CKMT1A | 0.384 | FGF1 | 1.751 |
| C1orf106 | 1.896 | CTSB | 9.211 | FXYD3 | 7.425 | FOXC2 | -3.978 |
| C4orf19 | 2.445 | EDNRA | 4.930 | HPGD | 6.467 | GADD45G | 1.786 |
| CDH1 | 8.627 | FOXC2 | 1.352 | MITF | 7.339 | INHBA | 1.962 |
| CEACAM1 | 4.592 | IFITM3 | 10.371 | MTUS1 | 3.385 | KRT16 | 2.625 |
| CGN | 5.918 | ITGA5 | 5.987 | MYH14 | 5.677 | KRT7 | 7.880 |
| CLDN4 | 8.052 | MMP3 | 3.602 |  |  | NID2 | 3.067 |
| CLDN7 | 7.632 | POSTN | 9.209 |  |  | NOTCH3 | 4.381 |
| CX3CR1 | 3.235 | SERPINE1 | 6.068 |  |  | PDGFRB | 5.254 |
| DSC2 | 4.658 | SPOCK1 | 4.503 |  |  | SERPINE1 | 2.100 |
| EHF | 6.185 | SULF1 | 6.699 |  |  | SPOCK1 | 1.611 |
| EPHA1 | 4.359 | TGFB1 | 6.076 |  |  | TPM2 | 4.213 |
| EXPH5 | 2.720 | WISP1 | 3.690 |  |  | VEGFC | 1.694 |
| FA2H | 2.587 |  |  |  |  | WISP1 | 3.202 |
| GPX2 | 1.897 |  |  |  |  | WNT5A | 2.836 |
| GRB7 | 5.585 |  |  |  |  |  |  |
| HPGD | 1.641 |  |  |  |  |  |  |
| ICA1 | 5.752 |  |  |  |  |  |  |
| IL20RA | 3.761 |  |  |  |  |  |  |
| IRF6 | 7.491 |  |  |  |  |  |  |
| JUP | 8.885 |  |  |  |  |  |  |
| MSX2 | 4.256 |  |  |  |  |  |  |
| POF1B | 2.093 |  |  |  |  |  |  |
| PPL | 5.551 |  |  |  |  |  |  |
| SH2D3A | 3.672 |  |  |  |  |  |  |
| TMPRSS2 | 3.378 |  |  |  |  |  |  |
| TUBBP5 | 1.727 |  |  |  |  |  |  |
| WNT3A | -2.375 |  |  |  |  |  |  |
| WNT4 | 2.885 |  |  |  |  |  |  |

**Fig. S1.**
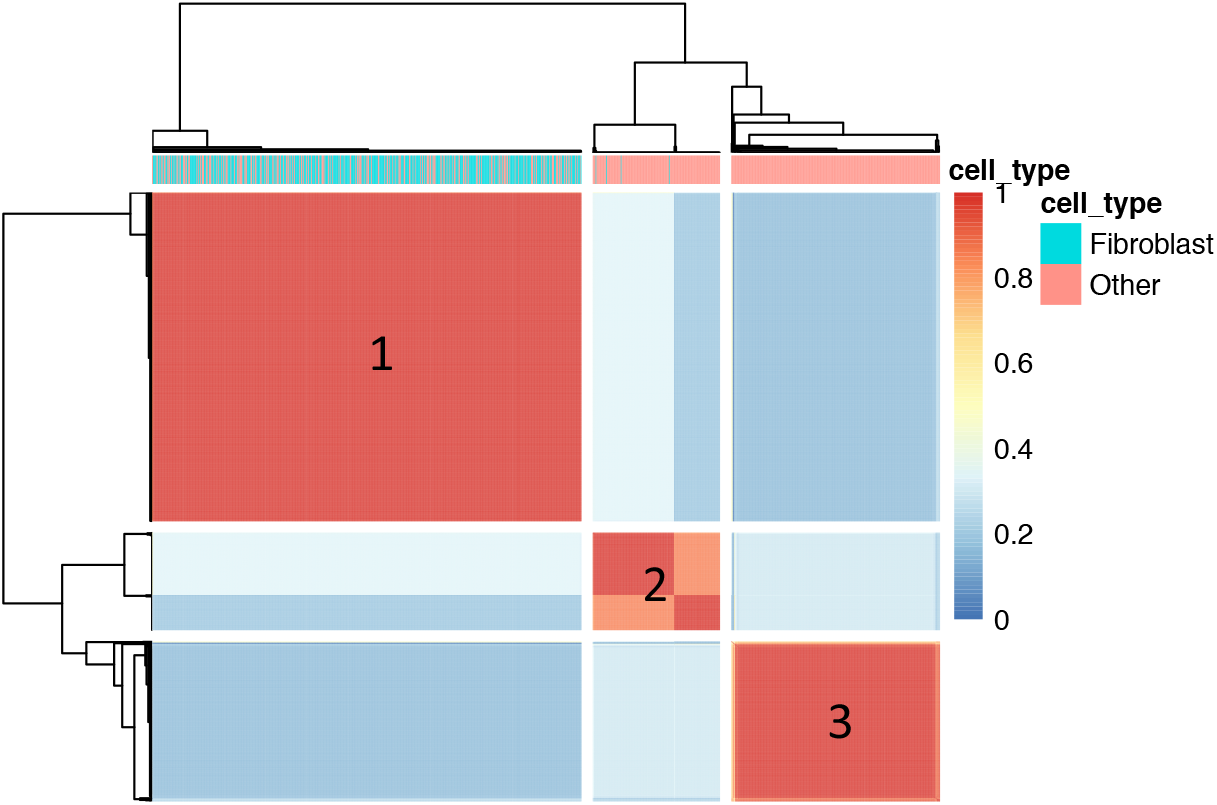
Consensus matrix for similarity and clustering of cell samples. The symmetric 1034×1034 matrix is colored in element(i,j) by similarity in assigning cells i and j to the same cluster when the clustering parameters are changed. A similarity score of 0 (blue) indicates that the two cells are always assigned to different clusters while a score of 1 (red) indicates that the two cells are always assigned to the same cluster. The similarity of the samples are also illustrated by the dendrograms shown on the top and side. The top bar indicates whether the cell was annotated as a fibroblast based on COL1A1 and COL1A2 expression (aqua - fibroblast, pink - other).

